# Molecular basis for GIGYF-TNRC6 complex assembly in miRNA-mediated translational repression

**DOI:** 10.1101/2021.08.20.457040

**Authors:** Meghna Sobti, Benjamin J. Mead, Cátia Igreja, Alastair G. Stewart, Mary Christie

## Abstract

The GIGYF proteins associate with 4EHP and RNA-associated proteins to elicit transcript-specific translational repression. However, the mechanism by which the GIGYF1/2-4EHP complex is recruited to its target transcripts remain unclear. Here we report the crystal structures of the GYF domains from GIGYF1 and GIGYF2 in complex with proline-rich sequences from miRISC-binding proteins TNRC6C and TNRC6A, respectively. The TNRC6 proline-rich motifs bind to a conserved array of aromatic residues on the surface of the GIGYF1/2 GYF domain, bridging 4EHP to Argonaute-miRNA mRNA targets. Our structures also reveal a phenylalanine residue conserved from yeast to human GYF domains that contributes to GIGYF2 thermostability. The molecular details we outline here are likely to be conserved between GIGYF1/2 and other RNA-binding proteins to elicit 4EHP-mediated repression in different biological contexts.

## Introduction

Cap-dependent translation initiation requires the assembly of eIF4F, a heterotrimeric complex comprising the RNA helicase eIF4A, the scaffold protein eIF4G and the cap-binding protein eIF4E to the mRNA 5’ m^7^GpppN cap structure. eIF4G not only bridges eIF4A and eIF4E but also recruits the preinitiation complex to capped mRNA, which in turn commences scanning and identification of the start codon (Sonenberg and Hinnebusch 2009; Jackson et al. 2010). Translation initiation can therefore be regulated by factors that modulate these interactions, such as the eIF4E-binding proteins (4E-BPs) that compete with eIF4G binding to eIF4E (Mader et al. 1995; Marcotrigiano et al. 1999), or proteins that recognise the 5’ cap structure which do not associate with eIF4G. One such cap-binding protein is 4EHP (eIF4E homologous protein, also known as eIF4E2), which prevents the assembly of the eIF4F complex on target mRNAs to prevent translation initiation (Rom et al. 1998; Joshi et al. 2004; Hernández et al. 2005). In contrast with eIF4E, 4EHP interacts with GIGYF1 (Grb10-interacting GYF protein 1) and its paralog GIGYF2 (Morita et al. 2012; Chapat et al. 2017), the interaction of which closely resembles the eIF4E:eIF4G and eIF4E:4E-BP complexes (Peter et al. 2017). Knockout of *4ehp* or *Gigyf2* in mice results in perinatal and early postnatal mortality, respectively (Giovannone et al. 2009; Morita et al. 2012). Different neurodegenerative presentations are also associated with the genetic insufficiency or loss of GIGYF1/2 in animals and humans including schizophrenia, autism spectrum disorders and age-related neurodegeneration (Giovannone et al. 2009; Iossifov et al. 2014; Krumm et al. 2015; Thyme et al. 2019; Satterstrom et al. 2020).

Unlike eIF4G which recruits the translation initiation machinery, human GIGYF proteins have been shown to associate with proteins involved with mRNA degradation (eg. the carbon catabolite repression-negative on TATA-less [CCR4-NOT] deadenylation complex; ZFP36/tristetraprolin [TTP]), translational repression (DEAD-box RNA helicase DDX6), mRNA decapping (DDX6, PatL1), ribosome quality control (eg. ZNF598), and miRNA-mediated silencing (eg. TNRC6 proteins) (Ash et al. 2010; Morita et al. 2012; Fu et al. 2016; Schopp et al. 2017; Amaya Ramirez et al. 2018; Peter et al. 2019; Ruscica et al. 2019; Tollenaere et al. 2019; Mayya et al. 2021). TNR6CA (also known as GW182), TNR6CB and TNRC6C are Argonaute-binding protein scaffolds that have been implicated in mRNA degradation and translational repression of miRNA targets (Jonas and Izaurralde 2015).

Mutational analysis of zebrafish TNRC6A (*Dr*TNRC6A) indicated that a highly conserved PPGL motif located within the TNRC6A silencing domain contributes to translational inhibition but does not affect deadenylation, the first step of mRNA degradation (Mishima et al. 2012). The PPGL motif is also present in the human TNRC6 proteins, as well as TNRC6/GW182 proteins in other organisms, suggesting that this motif might play a conserved role in TNRC6/GW182-mediated translational repression across metazoa (Mishima et al. 2012). A direct interaction has been observed between TNRC6A and a fragment of GIGYF2 that encompasses its central GYF domain (Schopp et al. 2017). Knockdown of GIGYF2 affects miRNA-mediated translational repression of an mRNA reporter, indicating that GIGYF2 is a regulator of Argonaute-miRNA (miRISC) activity (Schopp et al. 2017).

The human GIGYF proteins were named after their GYF adaptor domains which recognise proline-rich sequences (PRSs) conforming to the PPGΦ consensus (where Φ corresponds to a hydrophobic residue, except for tryptophan) (Kofler et al. 2005; Ash et al. 2010). The GYF domain takes its name from a conserved glycine-tyrosine-phenylalanine (GYF) motif that is part of the larger GFP-X_4_-[M/V/I]-X_2_-W-X_3_-GYF signature characteristic of the PRS-binding region. This hallmark of GYF domains forms a bulge-helix-bulge structural element that generates a hydrophobic ligand binding surface (Freund et al. 1999). GYF domains can be further divided into two subfamilies that are named after the proteins from which they were first identified: the splicing factor CD2BP2 (CD2 antigen cytoplasmic tail-binding protein 2) (Nishizawa et al. 1998), and *Saccharomyces cerevisiae* (*Sc*) Smy2 (Kofler et al. 2005) which is thought to be the GIGYF2 homolog in yeast. The two subfamilies are distinguished by the length of the β1-β2 loop, as well as by the residue located at position 8 of the GYF domain; a longer β1-β2 loop and a Trp at position 8 is characteristic of the CD2BP2 family, while a shorter β1-β2 loop and an Asp at position 8 typifies the Smy2 class of GYF domains (Kofler and Freund 2006). Only three GYF-domain containing proteins are encoded in the human genome, namely, CD2BP2, GIGYF1 and GIGYF2.

The specificity of 4EHP-mediated translational repression is thought to be imparted by the GYF domain of the GIGYF proteins, which serves as an adaptor to bridge RNA-binding proteins to 4EHP (Morita et al. 2012; Fu et al. 2016; Schopp et al. 2017; Weber et al. 2020). Mutations that prevent TTP from binding to the GIGYF2 GYF domain impair TTP-mediated repression in mammalian cells (Fu et al. 2016; Peter et al. 2019). Moreover, the repression activity of zebrafish TNRC6A/GW182 can be reduced by mutation of the conserved PPGL motif (Mishima et al. 2012). Mutations within the GIGYF GYF domains that prevent interaction with PPGΦ-containing sequences correspondingly disrupt their recruitment to endogenous partners (Weber et al. 2020). The importance of the GIGYF GYF domain is further illustrated in tethering assays whereby the isolated GIGYF2 GYF domain displays repressive activity comparable to that of the full-length protein (Amaya Ramirez et al. 2018).

To understand the mechanism by which PPGΦ containing proteins are recruited by the GIGYF:4EHP complex, we determined the crystal structures of GIGYF1 and GIGYF2 GYF domains in complex with PPGL peptides from TNRC6C and TNRC6A, respectively. These structures highlight the conserved molecular mechanism of PRS recognition by GYF domains and reveal a structural feature in metazoan GYF adaptors that contributes to domain stability.

## Results

### PPGΦ sequences can directly interact with the isolated GIGYF2 GYF domain

Recent work has detected a direct interaction between the C-terminal silencing domain of TNRC6C that encodes a PPGΦ motif (1369-1690; Fig. 1A) and a central portion of GIGYF2 that encompasses the GYF domain (residues 532-740) (Schopp et al. 2017). The interaction was diminished when the PPGL motif was mutated and when the GYF domain was deleted (residues 607-740 of GIGYF2) (Schopp et al. 2017). We therefore examined whether the GIGYF2 GYF domain (residues 529-597) could alone interact with the isolated TNRC6C PPGL motif (residues 1470-1480, Fig. 1B, blue dashed box). Using glutathione-S-transferase (GST)-tagged TNRC6C PPGL motif, a direct interaction was detected in pull-down assays with His_6_-tagged GIGYF2 GYF domain (Fig. 1C). Similarly, GST-tagged PPGL fragments of TNRC6A and TNRC6B (Fig. 1B) pulled down the His_6_-GIGYF2 GYF domain (Fig. 1C). This analysis is consistent with interactions detected in other studies (Schopp et al. 2017), and demonstrates that the isolated GIGYF2 GYF domain is sufficient to directly bind to PPGL motifs of the TNRC6 paralogs.

**Figure 1.**
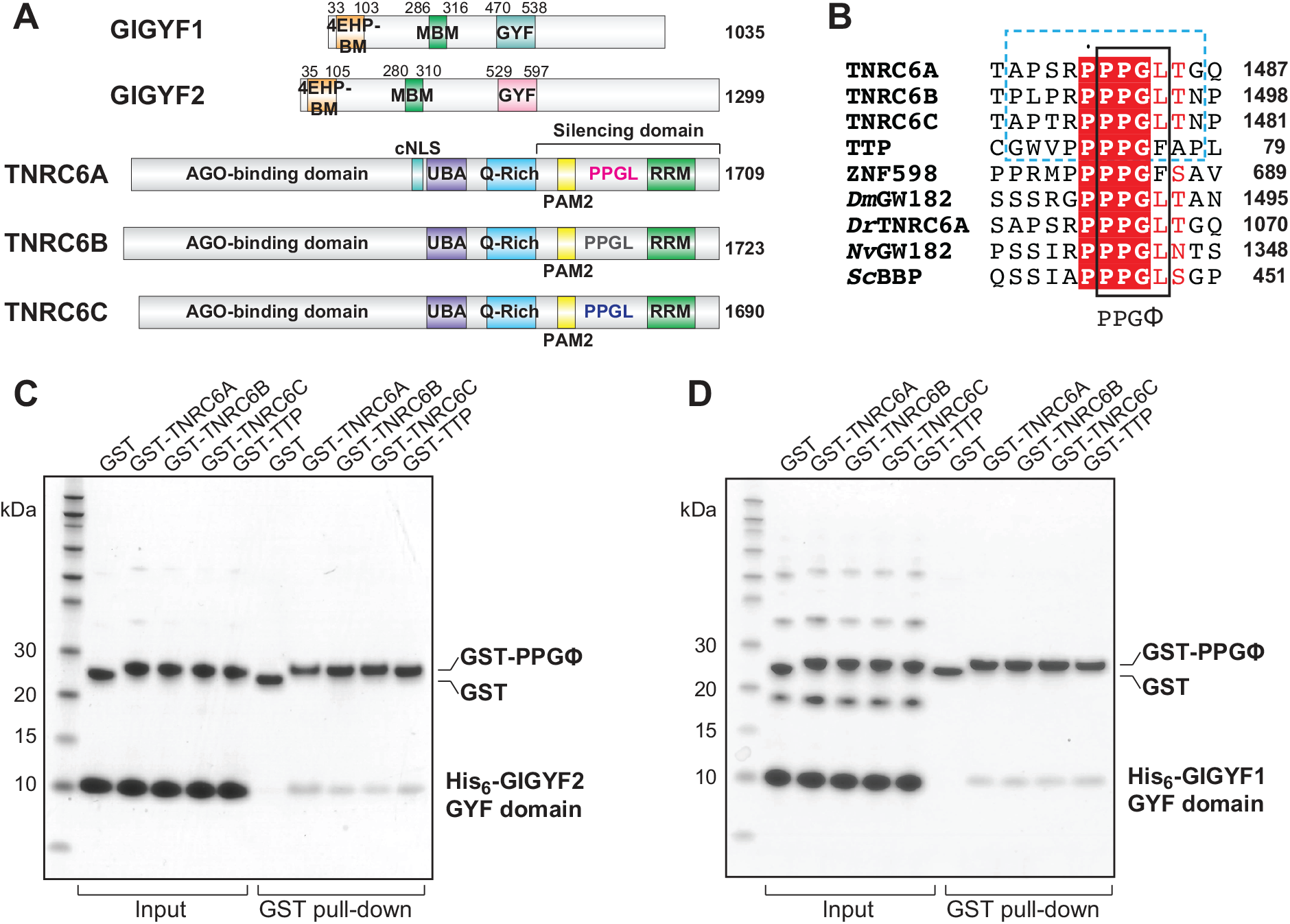
GIGYF1/2 GYF domains directly interact with proline-rich sequences from GW182/TNRC6 proteins. (A) Schematic representation of human GIGYF1/2 and TNRC6A-C proteins. GIGYF1/2 contain a 4EHP-binding motif (4EHP-BM) and a Me31B/DDX6-binding motif (MBM) within their N-terminal region, as well as a central GYF domain. The conserved PPGL motifs of the TNRC6 proteins are located within their C-terminal silencing domains, which also contains a PAM motif and an RNA-recognition motif (RRM) domain. TNRC6A contains a classical nuclear localisation signal (cNLS) between its Argonaute (AGO)-binding domain and ubiquitin-associated motif (UBA). The TNRC6 paralogs also contain Q-rich regions. (B) The PPGL motif is conserved in GW182/TNRC6A proteins in metazoa, and proline-rich sequences have been identified in RNA-binding proteins such as TTP and ZNF598. The PPGΦ is denoted by the black box, and the 11-residue peptides used in this study are indicated by the blue dashed box. For comparison, the PRS from *Sc*BBP is also shown. The species abbreviations are as follows: *Dm* (*Drosophila melanogaster*), *Dr* (*Danio rerio*), *Nv* (*Nematostella vectensis*), *Sc* (*Saccharomyces cerevisiae*). (C and D) GST pull-downs using recombinant His_6_-GIGYF1/2 GYF domain and GST-PRS sequences. GST only served as a negative control.

Direct interactions have also been observed between full-length GIGYF2 and TTP from mouse (Fu et al. 2016). We therefore tested whether this interaction was conserved in humans. A GST-tagged fragment of the first tetra-proline region of TTP (residues 68-78, Fig. 1B), which encodes a PPPPGF motif, could also interact with the GIGYF2 GYF domain (Fig. 1C).

### PPGΦ sequences can directly interact with the isolated GIGYF1 GYF domain

Co-immunoprecipitation assays using U2OS cells have detected an interaction between full-length GIGYF1 and TTP (Tollenaere et al. 2019). Deletion of the GYF domain or disruption of the characteristic ‘GYF’ motif in the context of the full-length GIGYF1 protein abolished its interaction with TTP, as well as with ZNF598 which contains three PPPPGF motifs (Tollenaere et al. 2019). Furthermore, a fragment encompassing the GYF domain of GIGYF1 (residues 260-540) was sufficient to bind to full-length ZNF598, and mutation of the three ZNF598 PPPPGF motifs could abrogate this interaction in U2OS cells (Tollenaere et al. 2019). As these results indicate that the GYF domain of GIGYF1 might interact with PPGΦ sequences in a similar manner to GIGYF2, we tested whether the isolated GIGYF1 GYF domain could also directly interact with the PPGΦ motifs from the TNRC6 proteins and TTP.

His_6_-tagged GIGYF1 GYF domain (residues 470-538; Fig. 1A) interacted with GST-tagged TNRC6 peptides (Fig. 1D). Similar to that observed for GIGYF2, the GST-tagged TTP peptide also directly interacted with the GIGYF1 GYF domain (Fig. 1D).

### The GIGYF1/2 GYF domains share structural characteristics typical of the Smy2 subclass of GYF adaptors

While molecular details are available for the CD2BP2 class of GYF domains in humans (Freund et al. 1999; Freund et al. 2002), no structural information is currently available for the human Smy2 class of GYF domains, namely the GYF domains from GIGYF1 and GIGYF2. We therefore determined the crystal structures of the isolated GYF domains from GIGYF2 and GIGYF1 in complex with PPGL peptides from TNRC6A and TNRC6C, respectively (Fig. 2A, 2B; Table S1). Crystals of the TNRC6C-GIGYF1 GYF domain complex were obtained in the C121 space group and diffracted to 1.79 Å resolution. The asymmetric unit comprises two TNRC6C-GIGYF1 complexes that display high similarity (rmsd of 0.4 Å over 61 Cα atoms). The TNRC6A-GIGYF2 GYF domain complex crystallized in the P2_1_2_1_2_1_ space group with one heterodimer in the asymmetric unit, and the structure was refined to 1.23 Å resolution.

**Figure 2.**
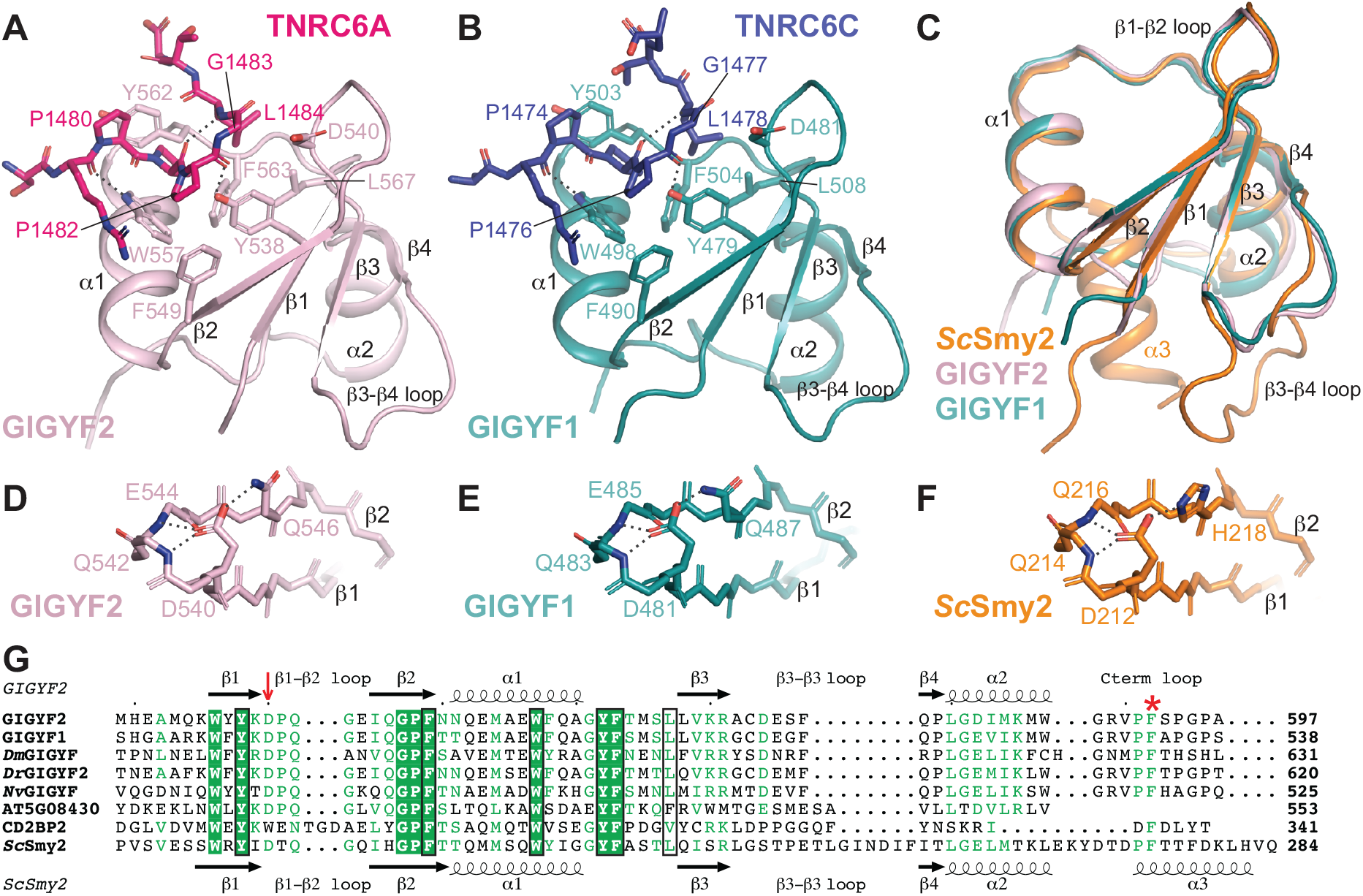
Overall structures of GIGYF1/2 GYF domains in complex with TNRC6 PPGL-containing peptides. (A) The TNRC6A PRS peptide (hot pink sticks) binds to a conserved arrange of aromatic residues on the surface of the GIGYF2 GYF domain (light pink cartoon). Hydrogen bonds are denoted by dashed lines. Similar interactions are observed between the TNRC6C PRS peptide (navy sticks) and the GIGYF1 GYF domain (teal cartoon) (B). (C) The GIGYF1/2 GYF domains are highly similar to the *Sc*Smy2 GYF fold (PDB ID 3FMA). (D-F) The conserved Asp residue that defines the Smy2 class of GYF domains mediates analogous interactions in GIGYF1/2 and *Sc*Smy2. (G) Structural-based sequence alignment of GYF domains. The prototypical Asp residue of the Smy2 GYFs is indicated by the red arrow. The residues that comprise the PRS-binding surface are boxed in black. The Phe plug is denoted by the red asterisk.

The GYF domains of GIGYF1 and GIGYF2 are comprised of a four-stranded antiparallel β-sheet with two α-helices that pack onto one face (Fig. 2A, 2B). The structures of the GYF domains from the human GIGYF proteins are highly conserved (Fig. 2C) with a backbone rmsd of 1.2 Å over 60 Cα atoms. The human GIGYF structures also display a high degree of structural similarity with *Sc*Smy2 GYF domain with rmsd values of 1.1 Å and 1.3 Å for GIGYF1 and GIGYF2, respectively (over 56 and 59 Cα positions). The secondary structures superimpose well, though there are some deviations in the β3-β4 loop, as well as the C-terminus of the domains whereby *Sc*Smy2 contains an additional α3 helix (Fig. 2C).

One of the defining features of the Smy2-subclass of GYF domains is a conserved Asp residue (D540 in GIGYF2, D481 in GIGYF1, and D212 in *Sc*Smy2; denoted by red arrow, Fig. 2G) within the β1-β2 loop (Kofler and Freund 2006; Ash et al. 2010). In our structures, GIGYF2 D540 and GIGYF1 D481 interact with a hydrogen bond donor present on the β2 strand (Q546 in GIGYF2, and Q487 in GIGYF1), which is analogous to H218 in *Sc*Smy2 (Fig. 2D-2F). The orientation of D540 and D481 in the human GIGYF proteins is further stabilised by hydrogen bonds with the backbone amides of the β1-β2 loop (Q483 and E485 in GIGYF1, and Q542 and E544 in GIGYF2; Fig. 2D, 2E). Identical interactions are observed for D212 in the *Sc*Smy2 structure (Fig. 2F).

### The GIGYF1 and GIGYF2 GYF domains interact with PPGL sequences

The crystals of human GIGYF GYF domains were obtained in complex with PPGL peptides from TNRC6A and TNRC6C, and the peptides could be built unambiguously into strong density located at the canonical PRS-binding surface (Fig. 2A, 2B, S1A and S1B). A series of residues in the GYF domains form a hydrophobic PRS-binding groove (Fig. 3A, 3B). These residues are Y538, F549, W557, Y562, F563, L567 in GIGYF2, and Y479, F490, W498, Y503, F504, L508 in GIGYF1 (Fig. 3A, 3B). Two hydrogen bonds are formed between GIGYF2 residues Y538 and W557 and the TNRC6A peptide backbone (Fig. 2A). Analogous interactions are observed in the GIGYF1-TNRC6C (Fig. 2B), *Sc*Smy2-BBP and CD2BP2-BD2 complex structures (Freund et al. 2002; Ash et al. 2010).

**Figure 3.**
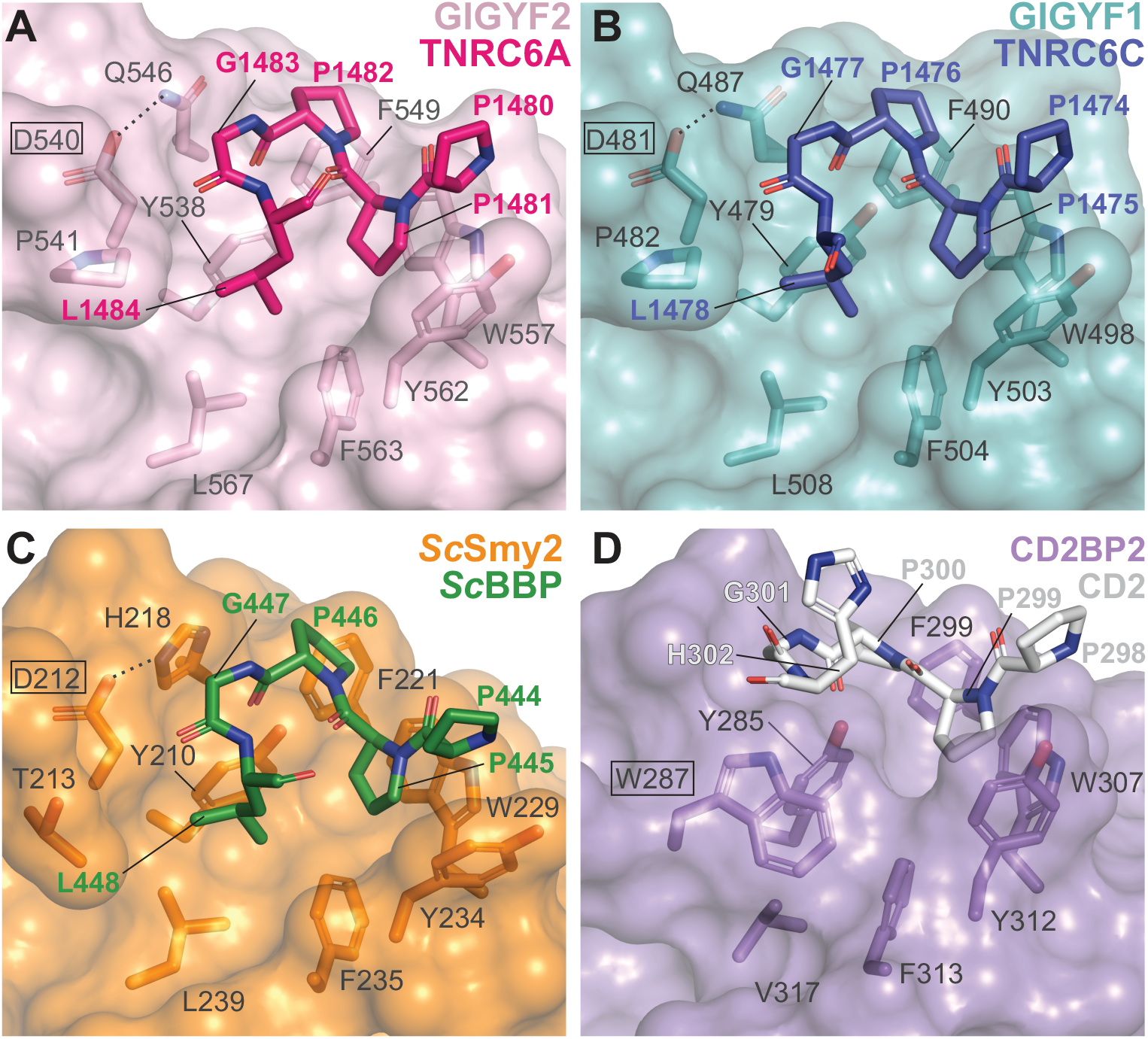
Proline-rich sequences bind to a conserved hydrophobic depression of the GYF domain surface. (A-C) Close-up views of TNRC6A, TNRC6C and *Sc*BBP peptides binding to the GYF domains of GIGYF2, GIGYF1 and *Sc*Smy2 (PDB ID 3FMA), respectively. The defining Asp residue of the Smy2 class of GYF domains are indicated by black boxes. By contrast, the CD2BP2 class of GYF domains contain a Trp (W287 in human CD2BP2, boxed; PDB ID 1L2Z) at this position (D). The conserved Trp in CD2BP2 GYF domains disrupt the hydrophobic cavity, which is exploited by PRS-containing sequences in the Smy2 class of GYF domains.

The TNRC6A PPGΦ motif adopts a polyproline II (PPII) helix with P1481 and P1482 interacting with the hydrophobic surface comprised of GIGYF2 Y538, F549, W557 and Y562. In this conformation, P1481 stacks against W557 in an aromatic pocket formed by F549 and Y562 (Fig. 3A). The TNRC6C PPGΦ motif similarly forms a PPII helical conformation, with P1475 stacking against W498 of GIGYF1 (Fig. 3B). The requirement for the Gly residue within the PPGΦ consensus is structurally rationalized as any larger side chain would clash with the conserved D540 and Q546 of the GIGYF2 GYF domain (or D481 and Q487 in GIGYF1, respectively) (Ash et al. 2010).

The Gly residue within the PPGΦ motif facilitates the adoption of turn in trajectory, enabling the subsequent hydrophobic residue (Φ) to insert into a hydrophobic cavity lined by Y538, F563 and L567 of GIGYF2 (Y479, F504 and L508 in GIGYF1). This cavity is surface exposed owing to the conformation of the characteristic Smy2-subclass Asp residue that orients away from this surface in GIGYF2, GIGYF1 and *Sc*Smy2 (boxed in Fig. 3A-3C). By contrast, the CD2BP2 subfamily of GYF domains contain a Trp in place of the Asp in the β1-β2 loop (Fig. 2G red arrow; boxed in Fig. 3D), which acts as a cover to obscure the hydrophobic surface formed by Y285, V317 and F313 (Fig. 3D) (Freund et al. 2002; Kofler and Freund 2006; Ash et al. 2010). The different surfaces created by the defining Asp and Trp residues in the Smy2 and CD2BP2 subfamilies, respectively, thereby determines the PRS binding specificities of the GYF domains by either extending or discontinuing the hydrophobic PRS binding surface (Ash et al. 2010).

TNRC6A and TNRC6C both contain Leu residues within their PPGΦ motifs, which are inserted into the Smy2 subclass-specific cavity of the GIGYF GYF domains. The TNRC6 peptides bind to the GYF adaptors in a manner highly similar to that observed when *Sc*BBP interacts with *Sc*Smy2 (Fig. 4C) (Ash et al. 2010). Our structures, together with the *Sc*Smy2-*Sc*BBP complex structure (Ash et al. 2010), reveal that the Smy2 subclass-specific cavity can accommodate larger hydrophobic sidechains, such as the Phe observed in the TTP and ZNF598 PPGΦ motifs (Fig. 1B). However, the cavity is too small to fit the Trp indole moiety without rearrangements within the binding pocket, consistent with the consensus identified by phage display analyses (Kofler et al. 2005).

**Figure 4.**
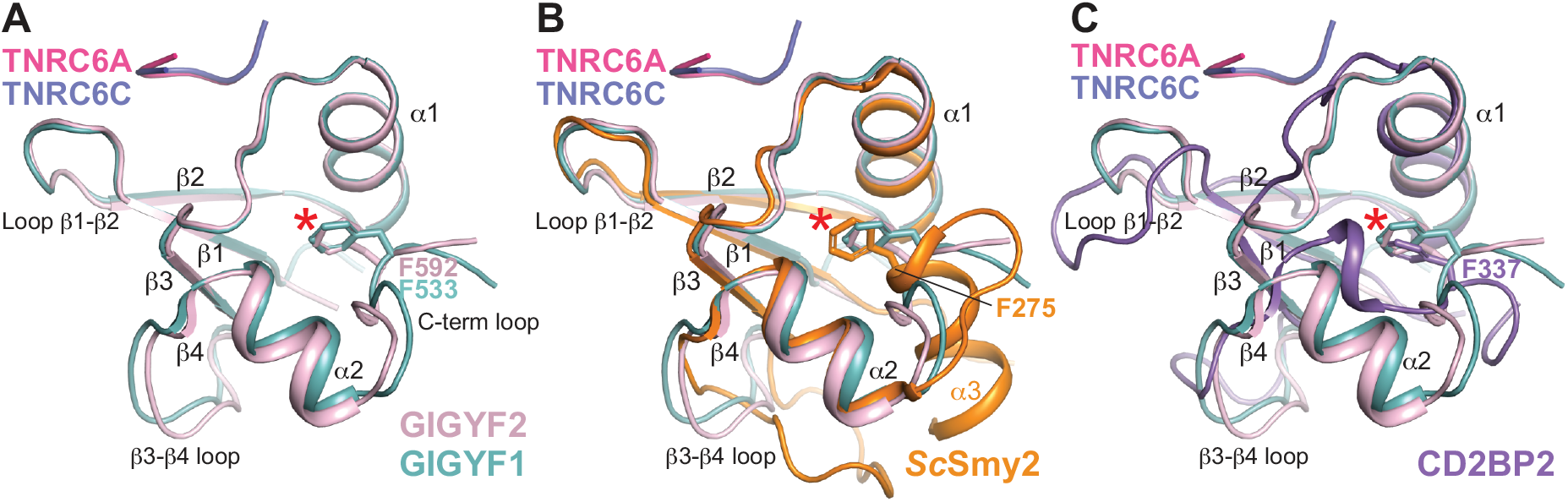
A conserved Phe residue is found at the C-terminus of Smy2 and CD2BP2 GYF domain classes from yeast to humans. (A) Superimposition of GYF domains from GIGYF1 and GIGYF2, shown in teal and light pink, respectively. The TNRC6 peptides are shown in hot pink and navy cartoon representation, The conserved Phe residue is shown in sticks and indicated by red asterisk. The same view is shown in (B) and (C) but with *Sc*Smy2 (orange; PDB ID 3FMA) and CD2BP2 (purple; PDB ID 1L2Z) superimposed, respectively.

### A conserved C-terminal Phe side chain is inserted into a hydrophobic pocket in GYF domains from yeast to humans

The N-terminal half of the GYF domains, comprising β1, β2, α1 and α1-β3 loop structural elements, are responsible for ligand binding and are highly conserved between GYF domain family members (Kofler and Freund 2006). By contrast, the C-terminal regions of GYF domains are poorly conserved, differing in length and composition (Kofler and Freund 2006). This is exemplified by *Sc*Smy2 which contains a C-terminal α3 helix (Ash et al. 2010) not present in the human GIGYF1/2 GYF domains (Fig. 2A-C, 2G). Due to the sequence identity between the GIGYF1 and GIGYF2 GYF domains (∼80% identical), even the C-terminal portions are highly similar with only slight differences observed in the conformations β3-β4 loop, and the C-terminal loop of the GYF domains (Fig. 4A).

Although there are slight differences in the C-terminal loop between the human GIGYF domain structures, the position of a Phe side chain adopts highly similar positions in GIGYF1 and GIGYF2 (F533 and F592, respectively; Fig. 4A denoted by red asterisk). Superimposition of the human GIGYF structures with *Sc*Smy2 reveals an analogous Phe side chain at this position encoded at the start of the α3 helix (Fig. 4B, red asterisk), despite the conformational variability observed between yeast and human proteins in this region. The structure of the *Sc*Smy2 GYF domain has previously been determined in a domain-swapped arrangement (PDB IDs 3K3V and 3FMA) with C-terminal α3 helices interacting with an adjacent GYF domain. Interestingly, the interactions between the F275 side chain are maintained in the domain-swapped conformation, which is inserted into the hydrophobic pocket of a neighbouring GYF domain (Ash et al. 2010). Moreover, a phenyl moiety is also observed in a similar orientation in the CD2BP2 GYF domain notwithstanding the large differences observed between the C-terminal regions of CD2BP2 and the Smy2-classes of GYF domains (Fig. 4C, red asterisk).

Close inspection of the C-terminal Phe residues in GIGYF1/2, *Sc*Smy2 and CD2BP2 (F592, F533, F275 and F337 in GIGYF2, GIGYF1, *Sc*Smy2 and CD2BP2, respectively) reveals that the Phe side chain is inserted, like a plug, into a hydrophobic cavity that is formed between the α1 and α2 helices and the inside face of the β-sheet (Fig. 5A-D). In this position, the conserved C-terminal Phe, which we term the ‘Phe plug’, forms part of the hydrophobic core of the small adaptor domains. Examination of GIGYF and CD2BP2 sequences from yeast to humans reveal strict conservation of this C-terminal Phe plug residue (Fig. 2G, and S2 denoted by a red asterisk) in both GIGYF and CD2BP2 GYF domain sequences. While this feature is not annotated as part of the GYF domain in some online databases, the general position of Phe plug was predicted by AlphaFold (Fig. S3A, S3B) (Jumper et al. 2021).

**Figure 5.**
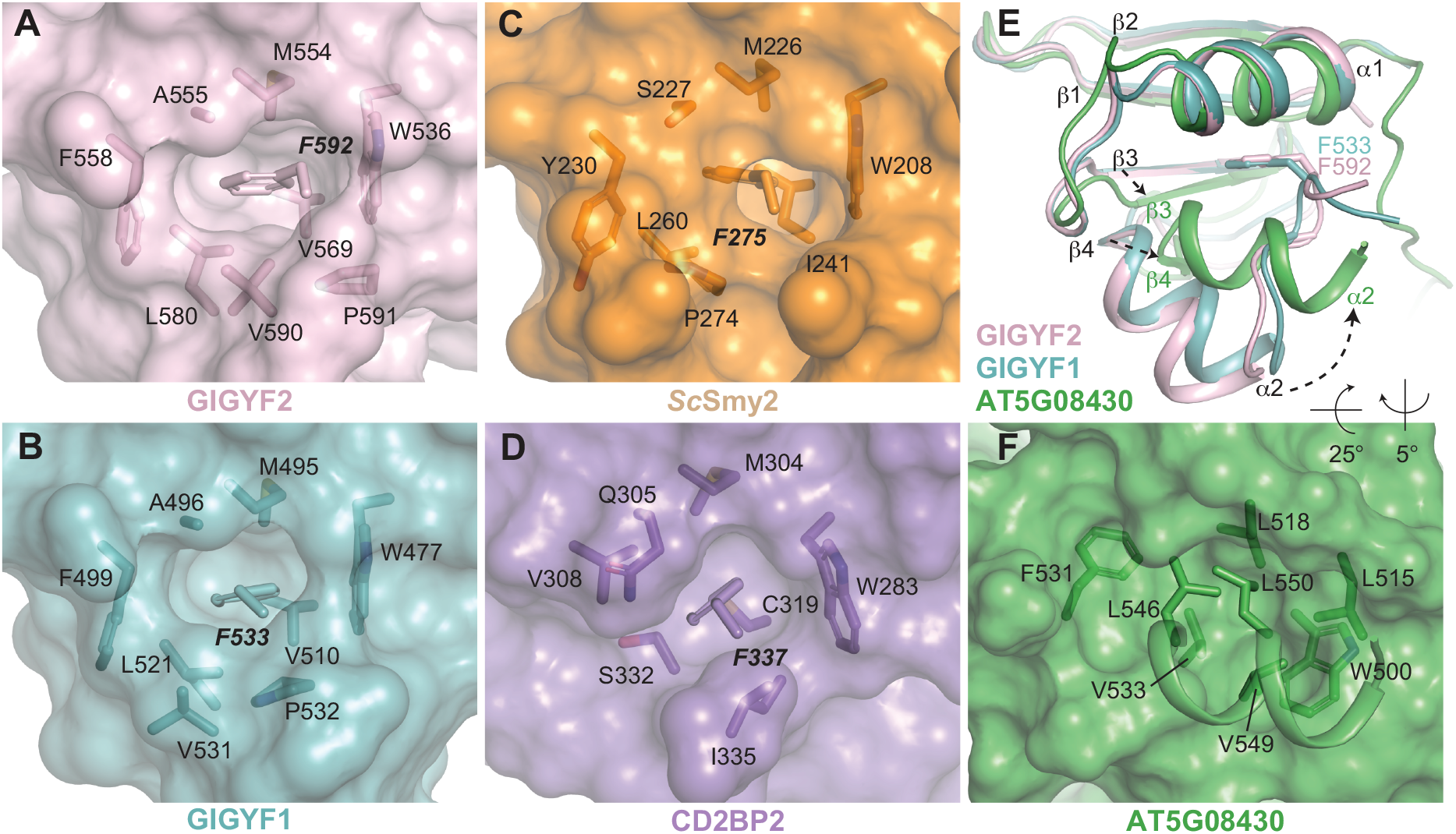
The conserved Phe plug forms part of a hydrophobic core in Smy2 and CD2BP2 classes of GYF domains from yeast to humans. (A to D) Surface representation of GIGYF2, GIGYF1, *Sc*Smy2 (PDB ID 3FMA) and CD2BP2 (PDB ID 1L2Z), respectively, with residues surround the Phe plug shown in sticks. (E) Superimposition of GIGYF2 and GIGYF1 (light pink and teal cartoon, respectively) with *A. thaliana* ATG08430 (green; PDB ID 1WH2). The GYF domain of *A. thaliana* ATG08430 is found at the extreme C-terminal end of the protein and does not encode a Phe plug residue. Conformational changes are observed between the *A. thaliana* ATG08430 and GIGYF1/2 in the α2 helix, as well as the β3 and β4 strands of the GYF domain. (F) The α2 helix in the plant GYF domain packs against the hydrophobic surface in the absence of a Phe plug.

### The Phe plug is not conserved in Arabidopsis GYF domains

By contrast, the Phe plug is absent in the only plant GYF domain who structure has been determined (from gene AT5G08430; Fig 5E, PDB code 1WH2). The *Arabidopsis thaliana* AT5G08430 GYF domain is more divergent from the human GIGYF structures than *Sc*Smy2 and CD2BP2 with larger RMSD values (2.4 Å and 2.3 Å for GIGYF1 and GIGYF2, respectively) and lower DALI server Z-scores (Z scores above 6.2 for CD2BP2 and *Sc*Smy2, and lower than 5.8 for AT5G08430).

The AT5G08430 GYF domain is located at the extreme C-terminal end of the protein with no additional residues encoded after the α2 helix (Fig. 2G). Compared with the α2 helix of GIGYF GYF domains, the AT5G08430 α2 helix is positioned closer to the α1 helix. In this orientation, hydrophobic side chains L546, V549 and L550 from the AT5G08430 α2 helix pack against the hydrophobic surface formed by W500, L515 and L518 (Fig. 5E-F). Assessment of other GYF or GYF-like domain sequences from *A. thaliana* identifies a Phe residue directly C-terminal to the α2 helix in three SWIB/PHD/GYF-domain containing proteins (AT2G18090, AT2G16485/NERD, AT3G51120; highlighted by dashed red box in Fig. S3C). However, other *A. thaliana* GYF domain-containing proteins (AT5G42950/EXA1, AT1G24300 and AT1G27430), for which no structures have yet been determined, do not encode obvious Phe plug residues at the C-terminus of their GYF domains. Notably, the GIGYF ortholog in *A. thaliana*, AT5G42950/EXA1, encodes a PPPGF sequence in this region that is thought to act as an autoinhibitory sequence to prevent low affinity interactions with the GYF domain (boxed blue in Fig. S3C) (Kofler and Freund 2006).

### The Phe plug contributes to GIGYF2 GYF domain stability

To investigate the importance of the conserved C-terminal Phe plug, we generated GYF domain mutants that disrupted the hydrophobic nature of the plug. More specifically, we replaced the Phe plug with the negatively-charged Glu residue generating GIGYF1 F592E and GIGYF2 F533E mutants. The His_6_-tagged mutant GYF domains were then tested for their ability to bind to the PRS sequences of TNRC6A and TTP. As the Phe residue is distal to the PRS binding site, the substitution did not affect the binding of PRS-containing sequences in GST pull-down assays under the conditions tested (Fig. S3D, S3E). Thermal stability assays, however, indicated that mutation of the Phe plug decreased the stability of the isolated GIGYF2 GYF domain with the apparent melting temperature of the F533E mutant 7 degrees lower than that observed for the wildtype domain (Tm_app_ of 61 °C and 54 °C for the wildtype and F533E GYF domains, respectively; Fig. S4).

## Discussion

In this study, we have elucidated the molecular basis of TNRC6 recognition by human GIGYF1/2 GYF domains, an interaction that enables 4EHP-mediated translational repression of miRNA-bound mRNA transcripts. While this study focuses on the TNRC6 proteins and TTP, other PRS-containing factors involved in translational regulation are known to interact with the GIGYF proteins in eukaryotic cells (Ash et al. 2010; Morita et al. 2012; Amaya Ramirez et al. 2018; Ruscica et al. 2019; Tollenaere et al. 2019; Mayya et al. 2021). As such, the molecular details we outline here are likely to be conserved in a variety of biological contexts to elicit 4EHP-mediated repression.

Using pull-down assays, we have demonstrated that the isolated GYF domains of GIGYF1 and GIGYF2 directly interact with PPGΦ-containing sequences from binding partners involved in post-transcriptional gene expression regulation, and that these interactions are conserved between the human GIGYF paralogs. This work extends previous studies by refining the minimal molecular requirements of the GIGYF-TNRC6/TTP interactions. Our structures reveal that the overall features of the Smy2 subfamily of GYF domains are conserved between yeast and humans, and rationalise how PPGF motifs, which are present in TTP and ZNF598, can bind to the GIGYF1/2 GYF domain surface. Comparison of our structures with reported GYF domains reveal the presence a conserved Phe plug at the C-terminus CD2BP2 and Smy2 subclasses of GYF domains from yeast to humans. Sequence analyses, however, indicate that this feature may not be present in *A. thaliana* GYF domains. Mutational analyses of the Phe plug in GIGYF1/2 suggests that this feature contributes to the stability of the GIGYF2 GYF domain, but the integrity of the Phe plug residue is not strictly required for PRS binding *in vitro*. This is consistent with the observation that the Phe plug is not present in the *A. thaliana* AT5G08430 GYF, although it is currently unclear if this domain can bind to PRS-containing proteins in plants. These results therefore refine our understanding of the molecular features of GYF domain adaptors, which should be taken into consideration when designing GYF domain constructs.

The “RPPPGL” sequence shared between TNRC6A and TNRC6C adopt similar conformations in our structures (Fig. 2A, 2B). It is therefore highly likely that TNRC6A interacts with GIGYF1 in a similar manner to that observed for TNRC6C, and, likewise, that TNRC6C interacts in an analogous manner to TNRC6A when binding to GIGYF2, corroborated by the similar binding properties seen in our pulldown studies (Fig. 1C, 1D). The “RPPPGL” motif is also present in TNRC6B (Figure 1B) (Mishima et al. 2012), and given the high similarity between the GIGYF2-TNRC6A and GIGYF1-TNRC6C structures, we would expect that comparable interactions would be mediated between TNRC6B and the GIGYF1/2 GYF domains. More broadly, the residues that comprise the GIGYF PRS-binding site, and the TNRC6/GW182 “PPGL” motif, are highly conserved from cnidaria (*Nematostella vectensis*; *Nv*) to humans (Fig. 1B, 2G). This indicates that the structures determined here would be suitable models of GIGYF-GW182 interactions in different species.

Mutations in GIGYF2 have been implicated in neurological conditions such as autism spectrum disorder, schizophrenia, and Parkinson’s disease (Lautier et al. 2008; Iossifov et al. 2014; Krumm et al. 2015; Thyme et al. 2019; Satterstrom et al. 2020). Although the association of GIGYF2 mutants in Parkinson’s disease remains controversial (Bras et al. 2009; Nichols et al. 2009; Tan and Schapira 2010), recent work has revealed the importance of the GIGYF2-4EHP axis in protein quality control (Hickey et al. 2020; Juszkiewicz et al. 2020; Sinha et al. 2020; Weber et al. 2020). It is therefore tempting to speculate that the impairment of GIGYF2 function may lead to the accumulation of misfolded and potentially cytotoxic polypeptide products that contribute to the development of neurological conditions. L580F is a GIGYF2 GYF domain mutation identified in Parkinson’s disease patient cohorts (Wang et al. 2010). As L580 forms part of the Phe plug binding pocket, the L580F mutation may therefore affect the packing of the GYF domain hydrophobic core. It will now be interesting to determine what effect the L580F mutation has on GYF domain stability and PRS-binding capacity.

## Materials and Methods

### Protein expression and purification

GST-PPGL and GST-TTP fragments were expressed in BL21(DE3) cells overnight at 18°C. The GST-fusion proteins were purified in 50 mM Tris 7.5, 125 mM NaCl, 2 mM 2-mercaptoethanol (BME). His_6_-GIGYF2 GYF domain was expressed in BL21(DE3) cells at 20°C overnight, and pellets were resuspended in 50 mM Tris 8.0, 400 mM NaCl, 2 mM imidazole, 5 mM BME. The His_6_-GIGYF2 GYF domain was eluted from a Ni-NTA column (GE Healthcare) using 250 mM imidazole and further purified using a S200 10/600 column (GE Healthcare) and flash frozen in 10 mM Tris 8.0, 150 mM NaCl, 5 mM BME. His_6_-GIGYF1 GYF domain was expressed and purified in similar conditions to GIGYF2. His_6_-GIGYF1/2 Phe plug mutants (F592E and F533E, respectively) were expressed and purified using similar conditions as the wildtype domains. Both wildtype and F592E mutant GIGYF1 GYF domains appeared to be less stable than the isolated GIGYF2 GYF domains. GIGYF2-TNRC6A complexes were obtained by immobilising GST-TNRC6A PPGL fragments on a GSTrap column and binding to purified GIGYF2 protein. The complex was eluted using 10 mM glutathione, and the GST tag was cleaved overnight using GST-3C protease. The complex was purified from GST using size exclusion chromatography. GIGYF1-TNRC6C complexes were purified using similar conditions.

### Crystallisation and data collection

Crystals of GIGYF2-TNRC6A were obtained in 1.0-1.4 M Na/K phosphate pH 7.4-7.8 and were flash frozen in liquid nitrogen using 25% glycerol as a cryoprotectant. Crystals of GIGYF1-TNRC6C were obtained in 100 mM Hepes 7.0, 1 M sodium malonate, and 25% glycerol was used as a cryoprotectant before freezing. Data were collected at the Australian Synchrotron MX2 beamline (McPhillips et al. 2002; Aragão et al. 2018) and processed using XDS (Kabsch 2010). For the GIGYF2-TNRC6A complex, *Sc*Smy2 was used as a molecular replacement model with *Sc*BBP coordinates removed (PDB ID 3FMA) (Ash et al. 2010). For the GIGYF1-TNRC6C, the GIGYF2 structure was used as the search model (TNRC6A coordinates removed) in PHASER (McCoy et al. 2007). Refinement was performed in Phenix (Liebschner et al. 2019), and both structures have excellent geometry (Table S1) (Williams et al. 2018).

### GST pull-down

Pull-down assays were performed essentially as described previously. Specifically, 80 μg of GST or GST-PPGL/GST-TTP fragments were incubated with 160 μg of His_6_-GIGYF2 or His_6_-GIGYF1 GYF domains in the presence of 25 μL of glutathione superflow agarose (Pierce) pre-equilibrated in binding buffer (50 mM Tris 7.5, 125 mM NaCl, 2 mM BME). The resin was washed 4 times in binding buffer, and bound proteins were eluted with binding buffer supplemented with 20 mM glutathione. The samples were analysed using SDS-PAGE, input lanes correspond to 2% of incubated protein and pull-down lanes correspond to 10% of eluted sample.

### Thermal shift assays

Purified His_6_-GIGYF2 GYF domain, wildtype and F533E mutant in 10 mM Tris 7.5, 150 mM NaCl, 5mM BME were mixed with SYPRO orange (Invitrogen; 10x final concentration) and dispensed into a 96-well qPCR plate. The solution was slowly heated from 20 ºC to 95 ºC using an Applied Biosystems real-time qPCR machine. Fluorescence was detected using a ROX filter. The data were analysed using the 7500 Software (ABI) and the minimum of the negative first derivative was used to determine the apparent melting temperature (Tm_app_). Assays were performed in triplicate. Similar calculations could not be performed with the isolated wildtype GIGYF1 GYF domain and the F592E mutant which both displayed high initial fluorescence and ambiguous melt transition, consistent with the instability of the domains observed during purification. Buffer alone and lysozyme was used as negative and positive controls, respectively.

## Data deposition

The coordinates of GIGYF1-TNRC6C (7RUQ) and GIGYF2-TNRC6A (7RUP) have been deposited into the Protein Data Bank.

## Acknowledgments

We thank the beamline scientists at the Australian Synchrotron for their assistance with data collection. This research was undertaken in part using the MX2 beamline at the Australian Synchrotron, part of ANSTO, and made use of the Australian Cancer Research Foundation (ACRF) detector. A.G.S. was supported by a National Health and Medical Research Council Fellowship APP1159347 and Grant APP1146403. M.C. was supported by Australian Research Council Fellowship DE160100608.

