## Supplementary material for "Molecular basis for GIGYF-TNRC6 complex assembly in miRNA-mediated translational repression"

**Table S1.** Data collection and refinement statistics

| | **GIGYF1-TNRC6C** | **GIGYF2-TNRC6A** |
|---|---|---|
| PDB code | 7RUQ | 7RUP |
| Space group | C121 | $P2_12_12_1$ |
| Unit cell | | |
| Dimensions a,b,c (Å) | 99.92, 32.50, 69.58 | 32.78, 38,31, 62.06 |
| Angles α,β,γ (°) | 90, 133.84, 90 | 90, 90, 90 |
| Data collection | | |
| Wavelength (Å) | 0.9537 | 0.9537 |
| Resolution | 29.63-1.79 | 19.43-1.23 |
| Observed reflections* | 99534 | 303634 |
| Unique reflections*# | 15377 (861) | 23420 (1130) |
| Completeness (%)*# | 99.5 (94.3) | 100 (100) |
| Multiplicity*# | 6.5 (5.9) | 13.0 (11.7) |
| $R_{merge}$ (%)*# | 9.6 (61.9) | 9.6 (129.7) |
| CC(1/2) *# | 0.997 (0.747) | 0.999 (0.778) |
| <I/σ(I)> *# | 11.2 (3.0) | 13.6 (2.0) |
| Wilson B ($Å^2$) | 19.6 | 11.5 |
| Refinement | | |
| Resolution | 29.63-1.79 | 19.43-1.23 |
| $R_{work}/R_{free}$ | 16.55/18.93 | 15.17/17.57 |
| Protein molecules/asu | 4 | 2 |
| Number of atoms | | |

| | | |
|---|---|---|
| Protein<br>Ligand/ion<br>Water | 2209<br><br>70 | 1148<br>5<br>96 |
| B-factors (Å$^2$)<br>Protein (Chains A,B,C,D)<br><br>Ligand/ion<br>Water | <br>24.7, 31.7, 27.0, 32.9<br><br><br>32.4 | <br>14.4, 18.3, -, -<br><br>24.0<br>26.5 |
| Ramachandran plot<br>Favored (%)^<br>Outliers (%)^ | <br>98.44<br>0 | <br>100<br>0 |
| Root mean square deviation<br>Bond lengths (Å)<br>Bond angles (°) | <br>0.011<br>1.238 | <br>0.012<br>1.276 |

*Output from Aimless

#Values in parenthesis are of the highest resolution shell

^Calculated by Molprobity

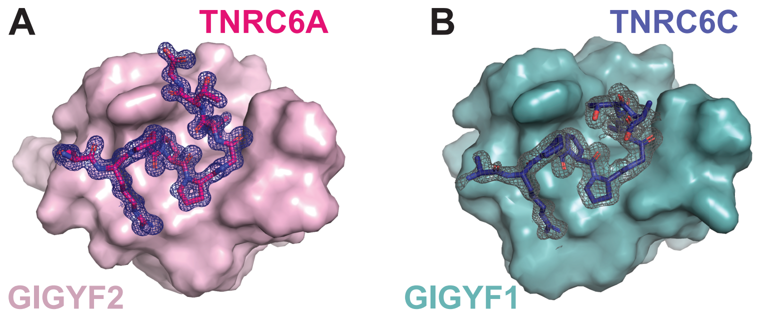


**Figure S1.** Structure of GIGYF1/2 GYF domains in complex with TNRC6 peptides. (A) The structure of GIGYF2-TNRC6A was refined to 1.23 Å and the final 2Fo-Fc density for the peptide is shown in blue mesh contoured at 1s. (B) The structure of GIGYF1-TNRC6C was refined to 1.79 Å and the final 2Fo-Fc density for the peptide is shown in grey mesh contoured at 1s.

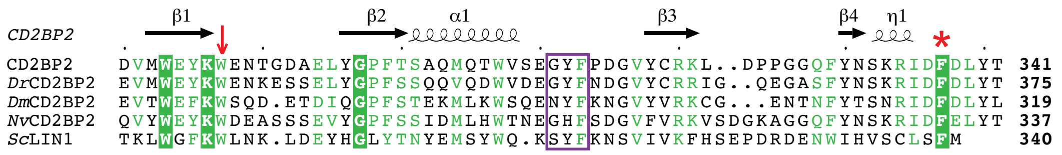

**Figure S2.** The Phe plug is conserved in CD2BP2 GYF domains from yeast to humans. Sequence alignment of CD2BP2 class GYF domains with the "GYF" motif boxed in purple. The conserved Phe plug residue is denoted by the red asterisk, and the defining Trp residue of the CD2BP2 class of GYF adaptors is indicated by the red arrow. The species abbreviations are as follows: *Dm* (*Drosophila melanogaster*), *Dr* (*Danio rerio*), *Nv* (*Nematostella vectensis*), *Sc* (*Saccharomyces cerevisiae*).

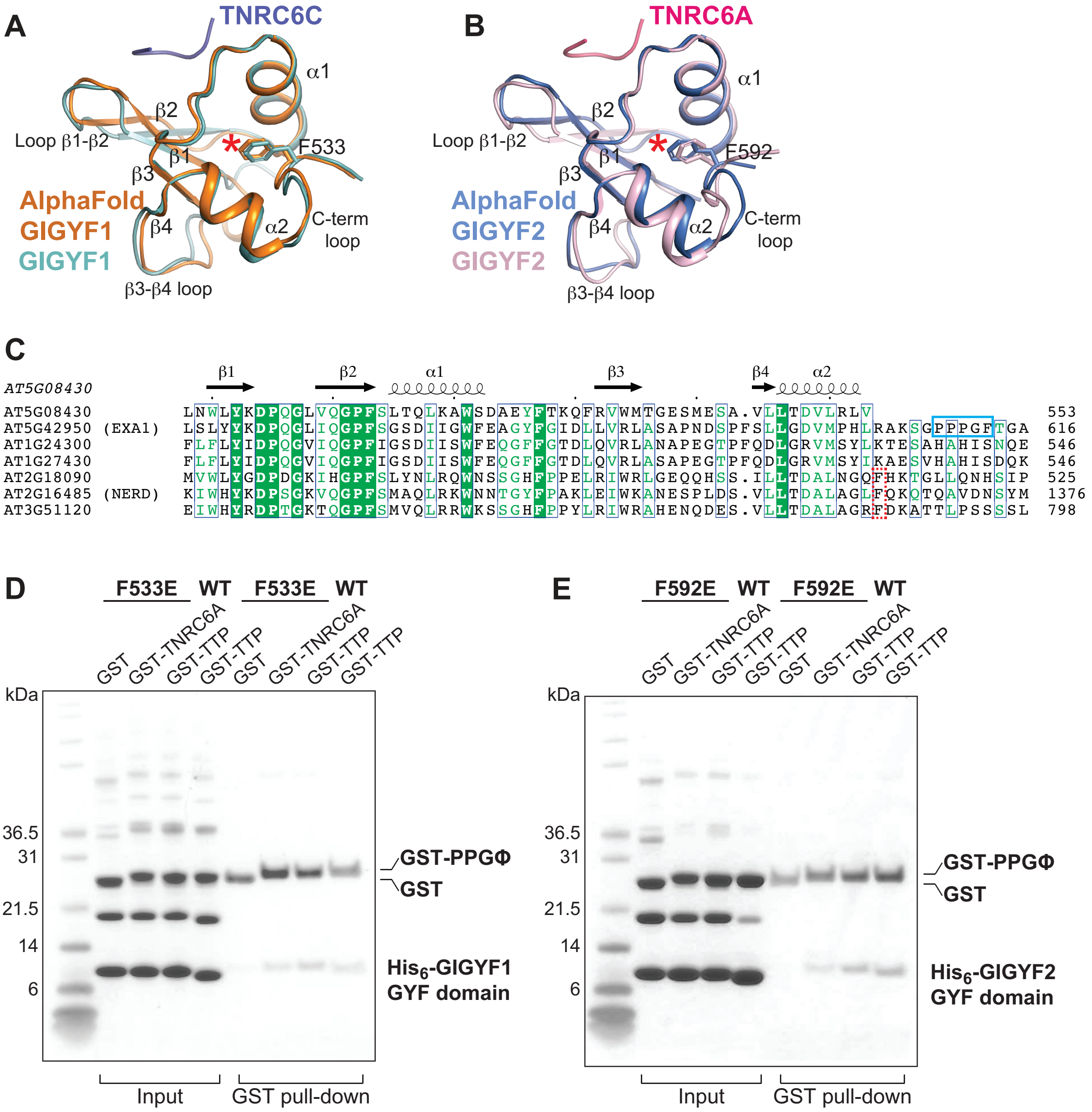


**Figure S3**. The integrity of the Phe plug is not strictly required for binding PRS peptides. (A and B) The position of the Phe plug was predicted by the AlphaFold (Jumper et al. 2021), validating the machine learning approach. (C) Alignment of GYF domains from *A. thaliana*. A Phe residue is located directly C-terminal to the α2 helix in some GYF domains (red dashed box). A "PPGF" motif is found at the C-terminus of the EXA1 GYF sequence (blue box). (D and E) Mutation of the Phe plug to a Glu does not prevent TNRC6 peptides from interacting with the GIGYF1/2 GYF domain. The wildtype $His_6$-GYF domains served as positive controls.

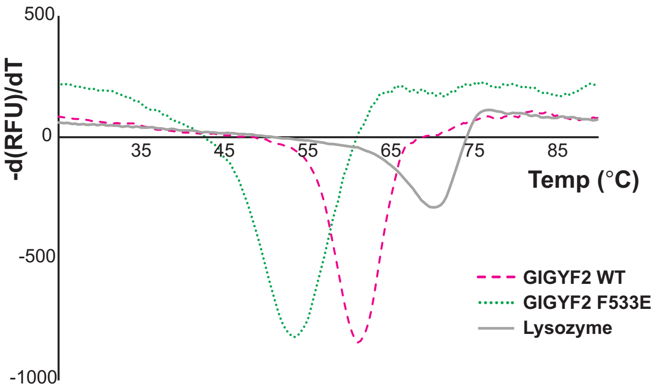


**Figure S4.** The Phe plug contributes to GIGYF2 GYF domain stability. Thermal shift assays using wildtype and F533E GIGYF2 GYF domains. Lysozyme was used as a positive control and the apparent melting temperature is similar to previous reports (Deore and Manderville 2019). Each assay was performed in triplicate and representative curves are shown.
